# Low-density Lipoprotein Receptor-related Protein 5 (LRP5)-deficient Rats Have Reduced Bone Mass and Abnormal Development of the Retinal Vasculature

**DOI:** 10.1101/2020.01.06.895797

**Authors:** John L. Ubels, Cassandra R. Diegel, Gabrielle E. Foxa, Nicole J. Ethen, Jonathan N. Lensing, Zachary B. Madaj, VARI Vivarium and Transgenics Core, Bart O. Williams

## Abstract

Humans carrying homozygous loss-of-function mutations in the Wnt co-receptor LRP5 (low-density lipoprotein receptor–related protein 5) develop osteoporosis and a defective retinal vasculature known as familial exudative vitreoretinopathy (FEVR) due to disruption of the Wnt signaling pathway. The purpose of this study was to use CRISPR/Cas9-mediated gene editing to create strains of Lrp5-deficient rats and to determine whether knockout of *Lrp5* resulted in phenotypes that model the bone and retina pathology in LRP5-deficient humans. Knockout of *Lrp*5 in rats produced low bone mass, decreased bone mineral density, and decreased bone size. The superficial retinal vasculature of Lrp5-deficient rats was sparse and disorganized, with extensive exudates and decreases in vascularized area, vessel length, and branch point density. This study showed that *Lrp*5 could be predictably knocked out in rats using CRISPR/Cas9, causing the expression of bone and retinal phenotypes that will be useful for studying the role of Wnt signaling in bone and retina development and for research on the treatment of osteoporosis and FEVR.

## Introduction

Wnt signaling plays key roles in development, and alterations in the Wnt pathway are among the most common events associated with human disease [1,2]. Wnts initiate cellular responses by binding to a receptor complex that includes a member of the Frizzled family of seven-transmembrane spanning receptors and either LRP5 or LRP6 (low-density lipoprotein receptor–related protein 5 or 6) [1]. Activation of this receptor complex inhibits the phosphorylation of the β-catenin protein by glycogen synthase kinase 3 (GSK3), which normally targets β-catenin for ubiquitin-dependent proteolysis. Thus, β-catenin is stabilized in the cytoplasm, and it can subsequently translocate to the nucleus and activate target gene transcription.

Almost 20 years ago, homozygous inactivating mutations in *Lrp5* were causally linked to the human syndrome osteoporosis pseudoglioma (OPPG) [3, 4]. Patients with OPPG develop osteoporosis in early childhood. Subsequent work, using genetically engineered mouse models, demonstrated that LRP5 most likely acts within the osteoblast lineage to regulate bone mass [5-9]. A large body of additional work further linked alterations in genes whose protein products interact with LRP5 (or with the highly homologous LRP6) to changes in human bone mass [1]. Primary among these products is sclerostin, a protein which is lost by genetic inactivation in patients with high bone mass [10].

OPPG patients have severely impaired vision at birth, associated with microphthalmia, retinal hypovascularization, and retrolental fibrovascular tissue (pseudoglioma) [11]. A related human hereditary disorder, familial exudative vitreoretinopathy (FEVR), can also be caused by inactivating mutations in *LRP5* [12]. In this case, progressive vision loss is due to decreased vascularization at the periphery of the retina [13]. This results in neovascularization in responseto oxygen deprivation and leads to retinal traction and subsequent retinal detachment. The new vessels are leaky due to an impaired blood–retina barrier, resulting in exudates. Mutations in other genes linked to the regulation of Wnt signaling, including Frizzled 4 and norrin, also cause FEVR and two related retinal vascular disorders, Norrie disease, and Coats disease [13,14].

Consistent with work in the skeletal system, genetically engineered mouse models have provided insights into the role of LRP5 in the development of the retinal vasculature. The normal development of the superficial vasculature is complete by postnatal day 8 and the deeper vasculature is complete by P21. In LRP5-null mice, vascular development is delayed, the vessels do not reach the periphery of the retina, and the deeper layers of the retinal vasculature do not develop. Neovascularization then occurs due to hypoxia, with the development of microaneurysms and vascular tufts [11, 15-17]. Fluorescein angiography reveals an impaired blood–retinal barrier [18].

The laboratory rat at one time was commonly used for physiologic studies, and genetic strains were developed through selective phenotypic breeding. The development of genetically engineered mouse models using embryonic stem cells led to the dominance of mice in biomedical studies, but these mouse models have some limitations. The relatively small size of mice confers a significant advantage in housing and husbandry costs, but while both mice and rats are rodents, functional differences exist. Further, it has been observed that genetically, mice and rats may be no more closely related than humans are to old world monkeys [19, 20]. On the other hand, a large physiologic data base suggests that in several fields, including learning and memory, neurologic function, and cardiovascular physiology, the rat is a better model for the study of human physiology and pathophysiology [19, 20]. Rats also have larger organs than mice, providing larger tissue samples and easier dissection and surgical manipuations.

The gene editting technology, CRISPR/Cas9, has allowed the reemergence of the rat as a key model organism in studying human disease, arguing for the expansion of genetically engineered rat models [21]. The development of genetically engineered mice using embryonic stem cell–based methods can take over 6 months. In contrast, CRISPR/Cas9 technology allows development of genetically modified rodents in only 2 months with greater predictability, and the ability to modify genes in zygotes makes it feasible to use rats for such studies. Already, genetically modified rats are being used in studies of neurophysiology and cardiovascular disease [22-25].

In the present study, we used CRISPR-Cas9-mediated methods to create three strains of rats carrying inactivating mutations in the second exon of the *Lrp5* gene. Studies were conducted to confirm knockout of the *Lrp5* gene and demonstrate lack of expression of functional LRP5 protein. The LRP5-deficient rats developed the low bone mass previously seen in LRP5-deficient humans and mice. The validity of the model is also supported by imaging and quantitative analysis of the retinal vasculature, showing that adult *Lrp5* knockout rats with impaired bone development also had a FEVR-like phenotype.

## Methods

### Experimental animals

Sprague-Dawley rats (Charles River Laboratories, Wilmington, MA) were maintained in accordance with institutional animal care and use guidelines, and experimental protocols were approved by the Institutional Animal Care and Use Committee of the Van Andel Institute.

### The generation of *Lrp5* knockout rats using CRISPR/Cas9

Three rat lines were created with a deletion in *Lrp5* by using a modified CRISPR/Cas9 protocol [26]. Two sgRNAs targeting exon 2 of *Lrp5* were designed using the MIT guide sequence generator (crispr.mit.edu.) against the rn5 genome reference sequence. The forward guide sequence, CCGCCGGGATGTACGACTAGTGG, and the reverse guide sequence, GTTGGCCACCTCGATGCGGTTGG, were cloned into vector pX330-U6-Chimeric_BB-CBh-hSpCas9, which was a gift from Feng Zhang (Addgene plasmid # 42230; http://n2t.net/addgene:42230; RRID:Addgene_42230). The T7 promoter was added to the sgRNA template, and the sequence was synthesized by Integrated DNA Technologies (Coralville, IA). The PCR-amplified T7-sgRNA product was used as template for *in vitro* transcription using the MEGAshortscript T7 kit (Thermo Fisher Scientific, Waltham, MA). The microinjection mix consisted of Cas9 mRNA (Sigma Aldrich, St. Louis, MO) (final concentration of 40 ng/μL) and sgRNA’s (20 ng/μL) in injection buffer (10 mM Tris; 0.1 mM EDTA, pH 7.5) and was injected into the pronucleus of Sprague-Dawley rat zygotes. After identifying founders we backcrossed the line to wild-type Sprague-Dawley rats twice before intercrossing to generate animals for our study.

### Genotyping by PCR-HMA and amplicon sequencing

Genomic DNA from rat tail biopsies was isolated by alkaline digestion. To genotype *Lrp5* knockout rats, we used the following primers—*Lrp5*-E2-Fwd (CCTCACCACTCCTGTTGTTT) and *Lrp5*-E2-Rev (CCTGCCAGAAGAGAACCTTAC)—to amplify a 354-bp product. Anticipating that the founders would have small deletions, a heteroduplex mobility assay [27] was used to define genotypes. Amplicons were subjected to denaturation–slow renaturation to facilitate formation of heteroduplexes using a thermocycler. Samples were resolved on polyacrylamide gels (10%), and mobility profiles were used to define genotypes. PCR using the genotyping primers above was performed to amplify indels for each founder for sequencing. The amplified products were cloned using the NEB PCR cloning kit (New England Biolabs, Ipswich, MA). Clones from each founder were Sanger-sequenced using the NEB analysis primers to define indels.

### Immunoblot analysis of LRP5 protein expression

The lack of expression of functional LRP5 protein was confirmed by immunnoblotting. Protein from rat tail biopsies was isolated in lysis buffer (50 mM Na_2_HPO_4_, 1 mM sodium pyrophosphate, 20 mM NaF, 2 mM EDTA, 2 mM EGTA, 1% Triton X-100, 0.5 mM DTT, protease inhibitor tablet Roche, Basal, Switzerland) using lysing matrix M (MP Biomedicals, Irvine, CA) for homogenization in the FastPrep-24 sample disruption instrument (MP Biomedicals). The samples were centrifuged at 21,000 x *g* for 30 min at 4□°C to pellet nonlysed tissue. We analyzed 40 μg of protein by SDS-polyacrylamide gel electrophoresis on an 8% Tris-glycine gel (Thermo Fisher Scientific) and transferred to PVDF western blotting membrane overnight at 4□°C. Immunoblotting was performed using the following antibodies: Lrp5 (D80F2) rabbit mAb (Cell Signaling, 5731), β-actin (13E5) rabbit mAb (Cell Signaling, 5125) (Cell Signalling Technology, Danvers, MA), and V5 Tag mAb, HRPP (Thermo Fisher Scientific, R96125).

### Plasmid construction

As will be shown, a gene product was expressed by the rat strain that had an 18-bp deletion in the *Lrp5* gene. To assess its potential signaling activity, site-directed mutagenesis was used to generate a human *LRP5* construct with the same 18-bp deletion. Using a previously generated human *LRP5*-V5 plasmid [28], we produced a human *LRP5*Δ131-136-V5 plasmid using the Q5 site-directed mutagenesis kit (New England Biolabs) with forward primer of (CTCAATGGCACATCCCGG) and a reverse primer of (GTTGGTCTCTGAGTCCGTC). The plasmid was sequenced to confirm the correct modification.

### Cell culture, transient transfections, and reporter studies

Twelve-well plates coated with poly-d-lysine were seeded with equal numbers of human HEK293-Super-TOPflash (STF) cells [29]. The cells were transfected by XtremeGene HP per the manufacturer’s specifications (Roche). When cells were about 80% confluent, a total of 500 ng of plasmid DNA was transfected per well. To measure the ability of constructs to activate a TCF-dependent reporter, 293-STF cells were co-transfected with 166 ng of plasmid to be tested, 166 ng of pCDNA3.1-lacZ plasmid, and empty vector DNA that was added to normalize the total amount of DNA in each transfection. Transfections were performed in duplicate. Cells were harvested 36 h after transfection by lysing in 100 μL of 1× reporter lysis buffer (Promega, Madison, WI). Lysates were pelleted, and 10 μL of each supernatant was added to 50 μL of luciferase assay substrate (Promega). A Synergy Neo HTS multimode microplate reader (BioTek, Winnoski, VT) was used to measure luciferase activity, and readings were normalized for transfection efficiency by measuring β-galactosidase activity as previously described [30]. Luciferase and β-galactosidase activity were measured in duplicate.

A linear mixed-effects model with random intercepts to account for repeated sampling via technical repeats was used to analyze these data via lmer in R v3.6.0 (https://cran.r-project.org/). Luciferase measures were log2-transformed to improve model fit based on normality of residuals. A plasmid directing the expression of B-galactosidase under the control of a CMV promoter was included as a fixed-effect to adjust for varying levels of success for the transfection. Linear contrasts with a Benjamini-Hochberg adjustment for multiple testing were used to test specific two-sided hypotheses of interest while maintaing a 5% false discovery rate. To check protein expression from the various expression constructs, the pellets were resuspended in 1× SDS sample buffer, boiled for 10 min, passed through a 26-gauge needle five times, and analyzed by SDS-PAGE followed by western blotting.

### Femoral bone mineral density analysis by DXA

Femoral aerial bone mineral density (BMD) of 6-month-old rats was measured by dual-energy X-ray absorptiometry (DXA) using a PIXImus II bone densitometer (GE Lunar) [5]. Rats were anesthetized via inhalation of 2% isoflurane (Henry Schein, Melville, NY) with oxygen (1.0 L/min) for 10 min prior to imaging and during the procedure (≤ 5 min). The right hindlimb was placed on a specimen tray in the densitometer for analysis. Bone mineral density was calculated by the PIXImus software based on the defined active bone area of the femur.

### Micro-computed tomography

The femur morphology of 6-month-old rats was evaluated using the SkyScan 1172 micro-computed tomography (µCT) system (Bruker MicroCT, Kontich, Belgium). Femurs from 6-month-old rats were isolated, fixed in 10% neutral buffered formalin at room temperature for 48 h, and transferred to 70% ethanol for storage. Femurs were scanned in a 70% ethanol solution using 2000 x 1200 pixel resolution and 7 µm image pixel size. An X-ray voltage of 70 kV, current of 142 µA, and 0.5 mm aluminum filter were used. 2D cross-sectional images of the femurs were reconstructed using the reconstruction software, NRecon 1.7.4.6 (Bruker MicroCT).

A volume of interest (VOI) was defined using DataViewer 1.5.6.3 (Bruker MicroCT), and a region of interest (ROI) was defined using CTAn 1.18.8.0 (Skyscan, Bruker). The ROI for trabecular bone was drawn in the distal epiphysis, with a defined height of 3.08 mm beginning 0.775 mm proximal from the growth plate. The ROI for cortical bone was 1.002 mm in height and began 7.002 mm from the growth plate. Average thresholds were calculated from trabecular and cortical grayscale images separately and were used to run 2D and 3D analysis for cortical bone and trabecular bone, respectively. Parameters measured for trabecular bone include bone mineral density (BMD), bone volume/tissue volume (BV/TV), trabecular thickness, trabecular separation, and trabecular number, while parameters measured for cortical bone include tissue mineral density (TMD), tissue area, bone area, cortical area fraction (bone area/tissue area), and cross-sectional thickness. Representative images were generated using CTvol 2.3.2.0 (Bruker MicroCT).

RStudio Version 1.1.453 was used to determine statistical significance for DEXA and microCT parameters. For percentage values, p-values were calculated using a beta linear regression model along with a least square means model. For all other values, p-values were calculated by using a generalized linear model along with a least square means model.

### Retinal fixation, staining, and imaging

Rats were humanely euthanized with CO_2_, the eyes were removed, and a 2-mm incision was made at the corneo-scleral limbus. The eyes were fixed in 4% paraformaldehyde in PBS for 1 h. The corneas and lenses were then removed and fixation continued for 3 h. Retinas were removed from the eyecups and incubated for 2 h in 1% Triton X-100 in PBS. Alexa Fluor 594– conjugated *Giffonia simplifolia* isolectin B_4_, having an excitation maximum at 594 nm and emission at 617 nm (Thermo Fisher Scientific), was dissolved in PBS containing 0.5 mM CaCl_2_. The retinas were incubated overnight at 4 °C in isolectin B_4_, which binds to the glycocalyx of vascular endothelial cells. The retinas were washed in PBS for 1.5 h, flat mounted on Superfrost slides (Fisher Scientific), and cover-slipped in Prolong Gold antifade reagent (Fisher Scientific) [15, 17].

Fluorescent images of the retinal vasculature were captured using a Zeiss Axio Imager A2 microscope, ET – DAPI/FITC/Texas Red filter (Carl Zeiss AG, Oberkochen, Germany), and BIOQUANT OSTEO 2019 v19.2.60 software (BIOQUANT Image Analysis Corp., Nashville, TN). The vascular networks were quantified using AngioTool software (http://angiotool.nci.nih.gov;) [31]. This open-source software measures vascular density (% vessels/total area), vessel length, and branching index (branch points/mm^2^). Data were analyzed by Kruskal-Wallis one-way ANOVA on ranks and Dunn’s method using SigmaPlot 12 (Systat Software, Inc., San Jose, CA).

## Results

Three independent rat lines carrying deletions in exon 2 of *Lrp5* were created by CRISPR/Cas9-mediated germline modification. After gene editing, one-cell-stage embryos were transferred to pseudopregnant females. A total of 15 pups were born, 3 of them containing indels within exon 2. These pups included a founder with an 18-bp deletion, designated *Lrp5*^*KO1*^, and a second founder with a 22-bp deletion at the sgRNA2 site, designated *Lrp5*^*KO2*^ (Fig. 1A). A third founder, designated Lrp5^KO3^, had an inversion coupled with small deletions in the exon at both the sgRNA1 (11 bp) and sgRNA2 sites (3 bp) (Fig. 1B). These three founder rats were then crossed with wild-type rats, resulting in germline transmission of each of the three modified alleles. Rats heterozygous for each of these mutations were intercrossed and tail tissue was collected for evaluation of amount of LRP5 protein from each of these three alleles. Consistent with the predicted frameshifts associated with *Lrp5*^*KO2*^ (22-bp deletion) and *Lrp5*^*KO3*^ (containing the inverted allele), we could not detect LRP5 protein in rats homozygous for these mutations. In contrast, *Lrp5*^*KO1*^, containing the 18-bp deletion, had detectable LRP5 protein (Fig. 1C). This deletion resulted in the loss of amino acids 131–136, which encodes the amino acid sequence RIEVAN within the first β-propeller (Fig. 2A).

**Fig. 1.**
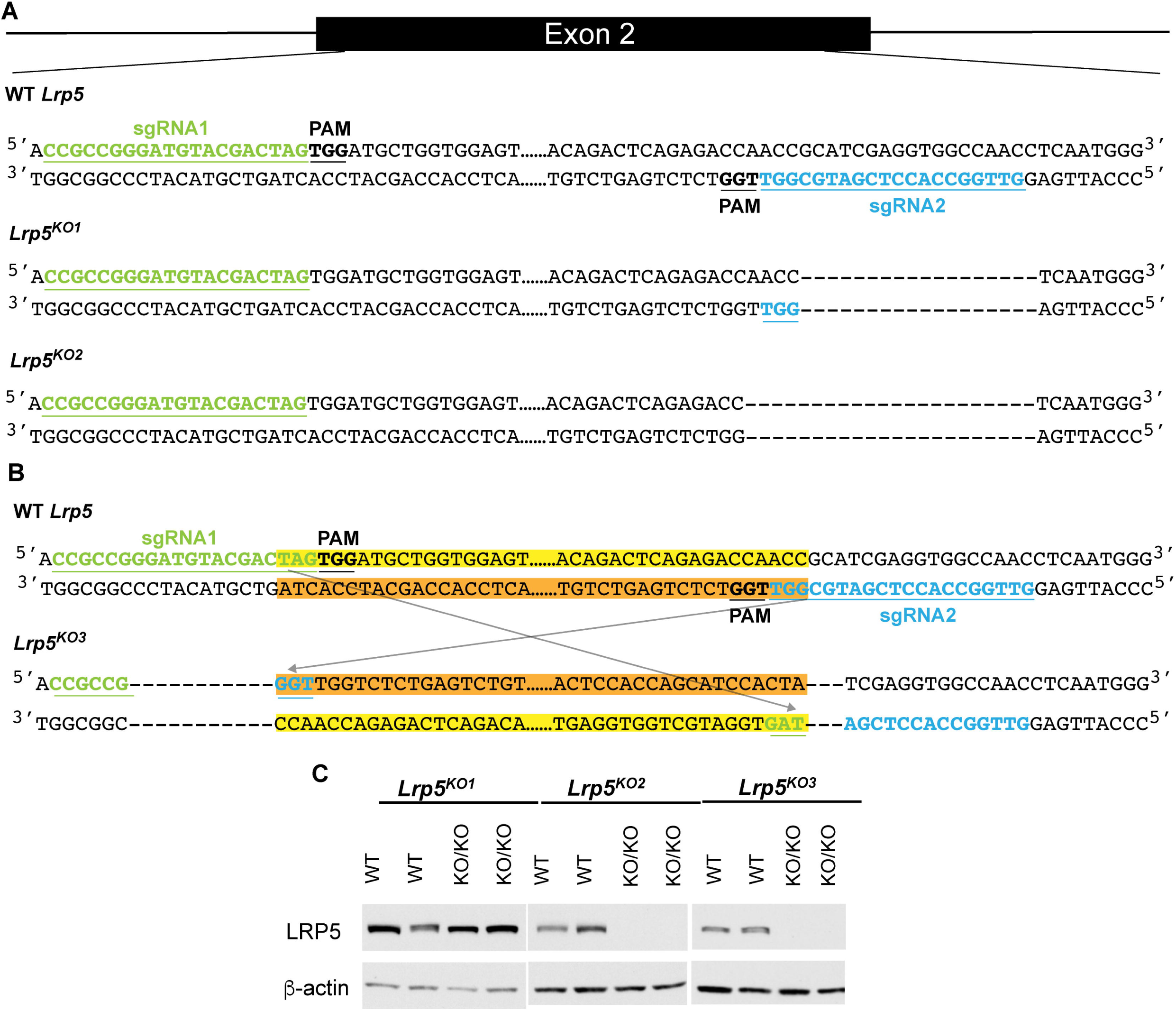
The *Lrp5*-deficient alleles created by CRISPR/Cas9-mediated genomic editing in the rat. **A.** Two sgRNAS (sgRNA1 and sgRNA2 with the PAM sequence noted) were injected with Cas9 into rat embryos to induce alterations within exon 2 of *Lr*p5. Two alleles with deletions were created and characterized: *Lrp5*^*KO1*^ had an 18-bp deletion (*LRP5* ^Δ131-136^) while *Lrp5*^*KO2*^ had a 22-bp deletion (indicated by dashes for each bp missing relative to the wild-type sequence). **B.** A third modified allele, *Lrp5*^*KO3*^, contained a modification caused by an inversion that likely occurred after Cas9-mediated cleavage at both sgRNA sites. Two small deletions associated with both sgRNA sites were also induced. **C.** Western blot analysis revealed a loss of Lrp5 protein from tissue lysates of rats homozygous for the *Lrp5*^*KO2*^ and *Lrp5*^*KO3*^ alleles. Rats homozygous for the *Lrp5*^*KO1*^ allele did express a protein (LRP5 ^Δ131-136^) from the modified allele.

**Fig. 2.**
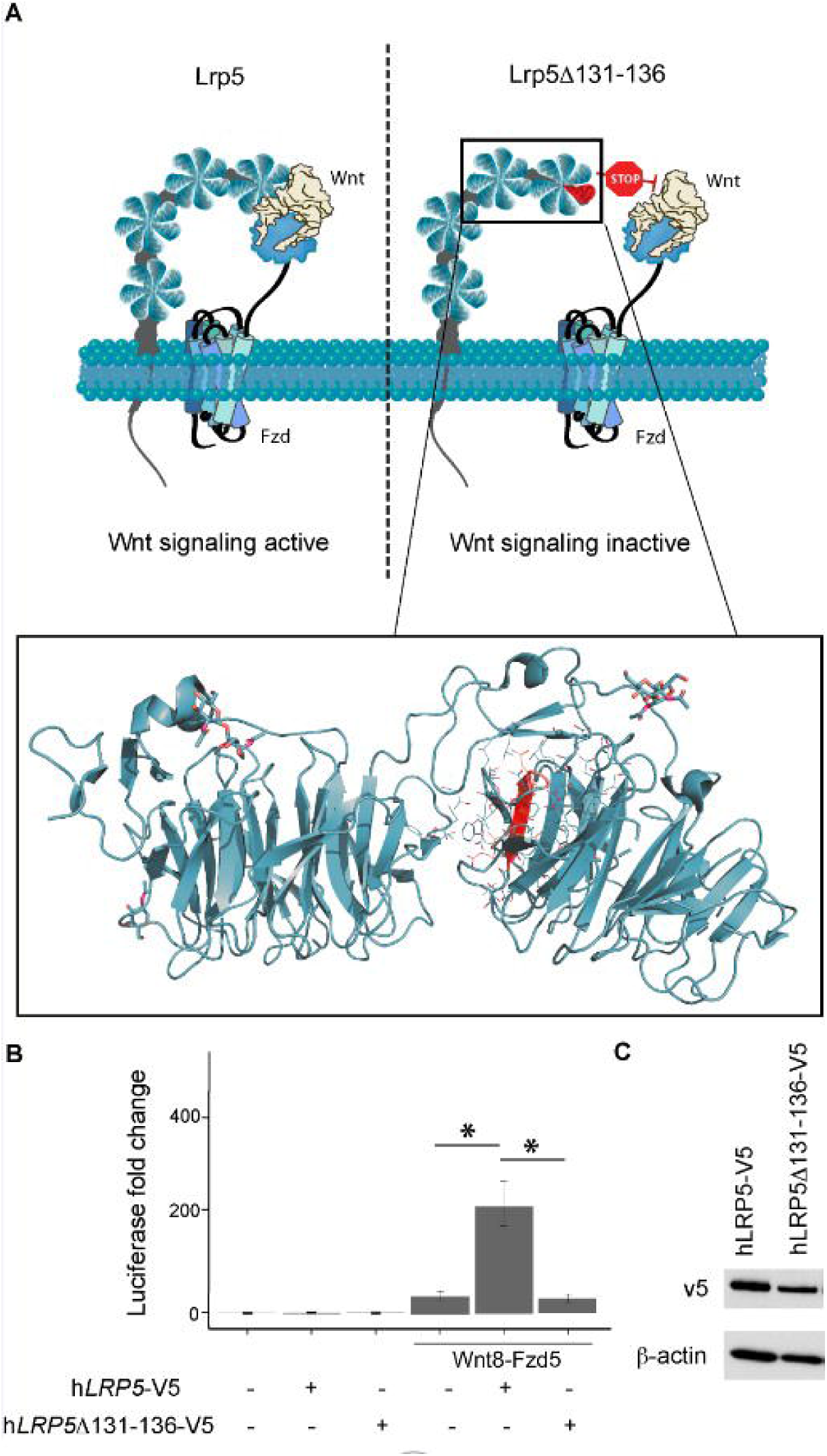
Functional characterization of the *Lrp5*^*KO1*^ protein, LRP5 ^Δ131-136^. **A.** Schematic of the Lrp5 protein showing the region (red) in the first β-propeller motif that contains the 18-bp in-frame deletion. **B** *LRP5* plasmids were co-transfected with a plasmid expressing the Wnt8-Fzd5 fusion protein into HEK293-STF Wnt/β-catenin reporter cells to define activation of the TOPflash luciferase reporter. Relative fold change of luciferase is shown (* = p < 0.05) **C.** The V5 epitope tag on the C terminus of both human *LRP5* expression plasmids was used to confirm protein expression. β-actin was used as a loading control.

To assess whether the in-frame deletion protein expressed in rats homozygous for the *Lrp5*^*KO1*^ mutation was active in Wnt/β-catenin signaling, we created a cDNA expressing human LRP5 ^Δ131-136^(hLRP5 ^Δ131-136^-V5) and measured its ability to activate a stably integrated β-catenin-responsive reporter gene (Super-TOPflash) in HEK293 cells [33]. We found that although transfecting an expression vector directing a wild-type version of human LRP5 with a V5 epitope (hLRP5-V5) could activate the TOPflash reporter in conjunction with an Wnt8-Fzd5 fusion protein [28], a plasmid directing LRP5 ^Δ131-136^ expression could not do so (Fig. 2B). We performed a western blot to validate qualitatively equivalent expression of the V5 tagged plasmids (Fig. 2C) This is consistent with the interpretation that the LRP5 ^Δ131-136^ protein expressed in *Lrp5*^*KO1*^ rats is not capable of supporting Wnt signaling.

### Effect of *Lrp5* knockout on bone

Initially, DXA was performed to investigate the skeletal phenotype in *Lrp5* knockout rats. This quantification showed aerial BMD of 6-month-old male and female femurs was significantly decreased in the *Lrp5*^*KO1*^, *Lrp5*^*KO2*^, and *Lrp5*^*KO3*^ rats compared to wild type animals (Suppl. 1). To further examine the skeletal phenotypes of *Lrp5* knockout rats, femurs from the *Lrp5*^*KO1*^, *Lrp5*^*KO2*^, and *Lrp5*^*KO3*^ lines were analyzed using micro-computed tomography (µCT). Both sexes in all three lines displayed the same trends in trabecular bone, but with varying severities (Fig. 3A). In the three female and male *Lrp5* knockout rat lines, we found significant decreases in bone mineral density (BMD; by 29-81%) and bone volume/tissue volume (BV/TV; by 84-92%) relative to wild-type controls (Fig. 3B-D). The decrease in bone mass was coupled with changes in trabecular morphology. The thickness of the trabeculae decreased significantly by 23-93% in all *Lrp5* knockout rats, resulting in greater separation. This increase in trabecular separation was significant in all *Lrp5* knockout rats with the exception of *Lrp5*^*KO2*^ female rats. The overall number of trabeculae in the distal femur of *Lrp5*^*KO1*^, *Lrp*^*KO2*^, and *Lrp5*^*KO3*^ rats decreased by 31-96%, 33-40%, and 44-48%, respectively (Fig. 3B-D). However, the decrease was only stastistically significant in *Lrp5*^*KO1*^ and *Lrp5*^*KO3*^ rats. Thus, the Lrp5 deficiency in *Lrp*^*KO1*^, *Lrp5*^*KO2*^, and *Lrp*^*KO3*^ rats negatively affected the trabecular bone framework.

**Fig 3.**
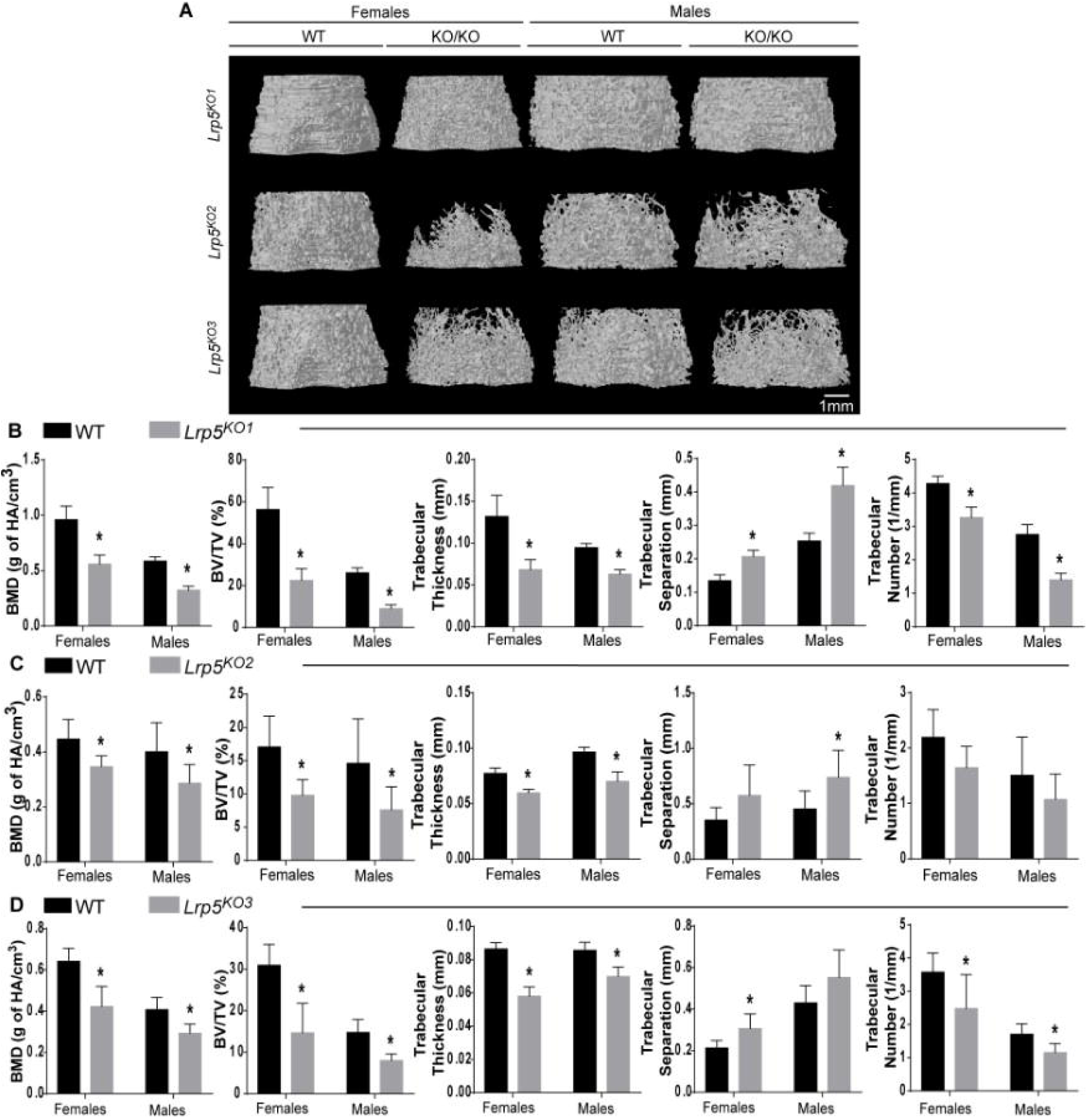
*Lrp5*-deficient rats showed a decrease in trabecular bone mass and quality. **A.** Representative µCT images of the trabecular bone region analyzed from the distal femur of *Lrp5*^*KO1*^, *Lrp5*^*KO2*^, and *Lrp5*^*KO3*^ rats. **B.** Quantitative analysis of bone parameters of *Lrp5*^*KO1*^ rats; bone mineral density (BMD), bone volume/tissue volume (BV/TV), trabecular thickness, trabecular separation, and trabecular number, *n* = 4-6. **C.** Quantitative analysis of the same bone parameters of *Lrp5*^*KO2*^ rats, *n* = 5. **D.** Quantitative analysis of the same bone parameters of *Lrp5*^*KO3*^ rats, *n =* 4-6. For all panels, * = *p* < 0.05.

Cortical bone architecture was also modified in both sexes (Fig. 4A). Cortical bone in *Lrp5* knockout rats was thinner and the size of femoral midshaft was smaller compared to the wild type animals.The cortical bone parameters were quantified by measuring tissue mineral density (TMD), cortical area fraction (CAF), tissue area, bone area, and cross-sectional thickness in *Lrp5*^*KO1*^, *Lrp5*^*KO2*^, and *Lrp5*^*KO3*^ rats (Figure 4B-D). TMD was significantly increased in *Lrp5*^*KO2*^ male and female rats alone, with no significant change in *Lrp5*^*KO1*^ or *Lrp5*^*KO3*^ rats compared to their wild type controls. Furthermore, all *Lrp5* knockout rats displayed no significant change in CAF relative to wild type rats. However, the change in cortical bone size was supported by the significant decrease in tissue area, bone area, and cross-sectional thickness of *Lrp5*^*KO1*^, *Lrp5*^*KO2*^, and *Lrp5*^*KO3*^ animals. The tissue area decrease in mutants was 14-42% in females and 18-34% in males, while the bone area decrease was 15-36% in females and 19-31% in males. Cross-sectional thickness significantly decreased by 6-14% in all *Lrp5* knockout rats with the exception of *Lrp*^*KO1*^ females. Overall, LRP5 deficency in mutant rats did not produce a consistent change in cortical bone density, but did result in an overall change in cortical structure.

**Fig 4.**
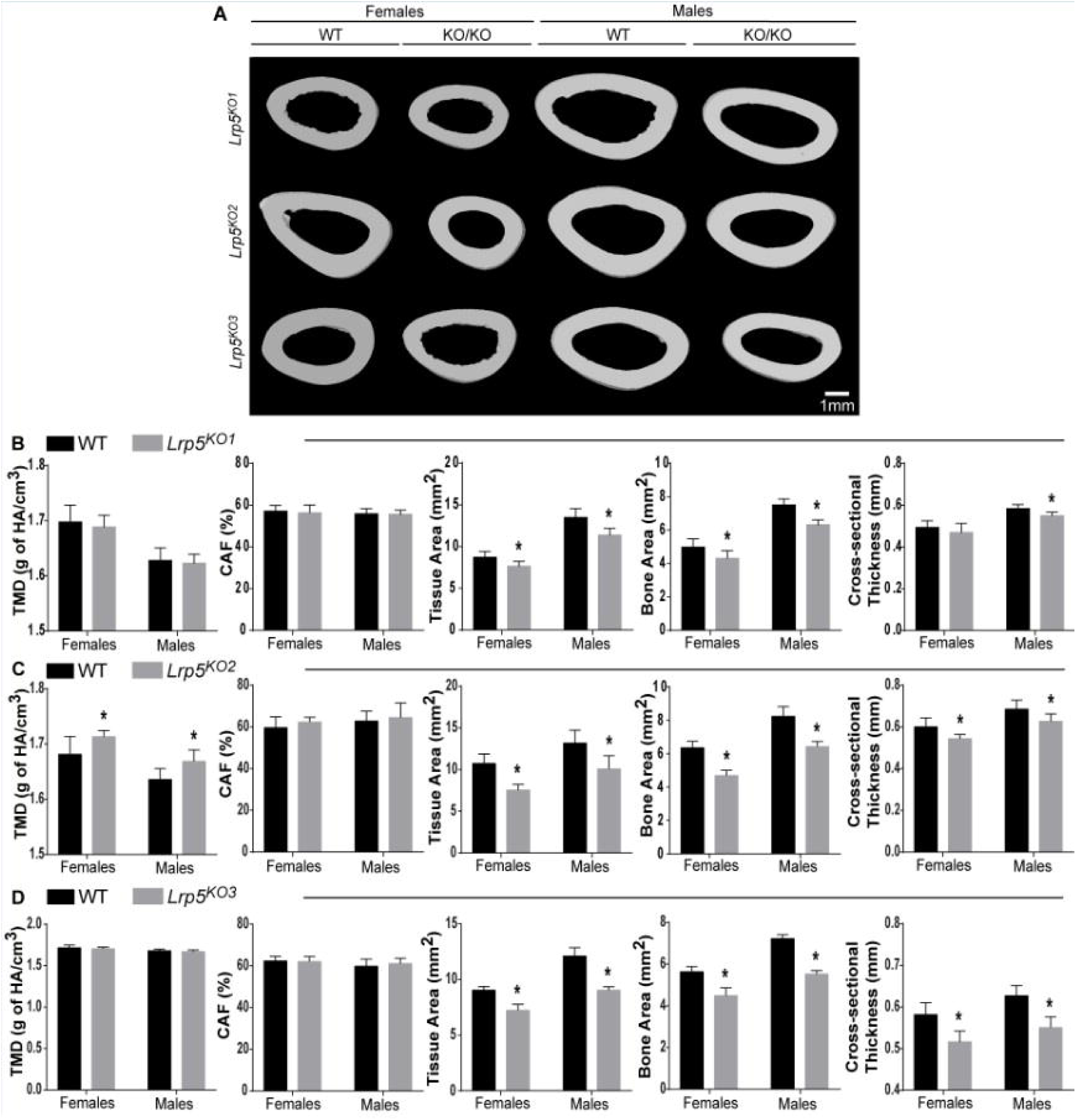
*Lrp5*-deficient rats showed a decrease in femoral cortical bone size. **A.** Representative µCT images of the cortical bone region analyzed from femurs of *Lrp5*^*KO1*^, *Lrp5*^*KO2*^, and *Lrp5*^*KO3*^ rats. **B.** Quantitative analysis of bone parameters of *Lrp5*^*KO1*^ rats; tissue mineral density (TMD), cortical area fraction (CAF, bone area/tissue area), tissue area, bone area, and average cross-sectional thickness, *n* = 4-6. **C.** Quantitative analysis of the same bone parameters of *Lrp5*^*KO2*^ rats, *n* = 5. **D.** Quantitative analysis of the same bone parameters of *Lrp5*^*KO3*^ rats, *n* = 4-6. For all panels, * = p < 0.05.

### Effect of *Lrp5* knockout on retinal vasculature

The retinal vasculature consists of three vascular beds: a superficial plexus in the nerve fiber layer of the retina, which was analyzed in this study; an intermediate network located in the inner plexiform layer; and a deep network in the outer plexiform layer [13]. These parallel beds originate at the optic disc from branches of the central retinal artery and return via the central retinal vein. Ten to twelve large superficial arteries and veins radiate from the optic disc of the rat retina, dividing to form the superficial network, as seen in the images of wild-type and heterozygous retinas in Figures 5 and 6. In contrast to bone, there was no difference in the retinas of male and female rats; therefore, the data from the sexes were combined.

**Fig. 5.**
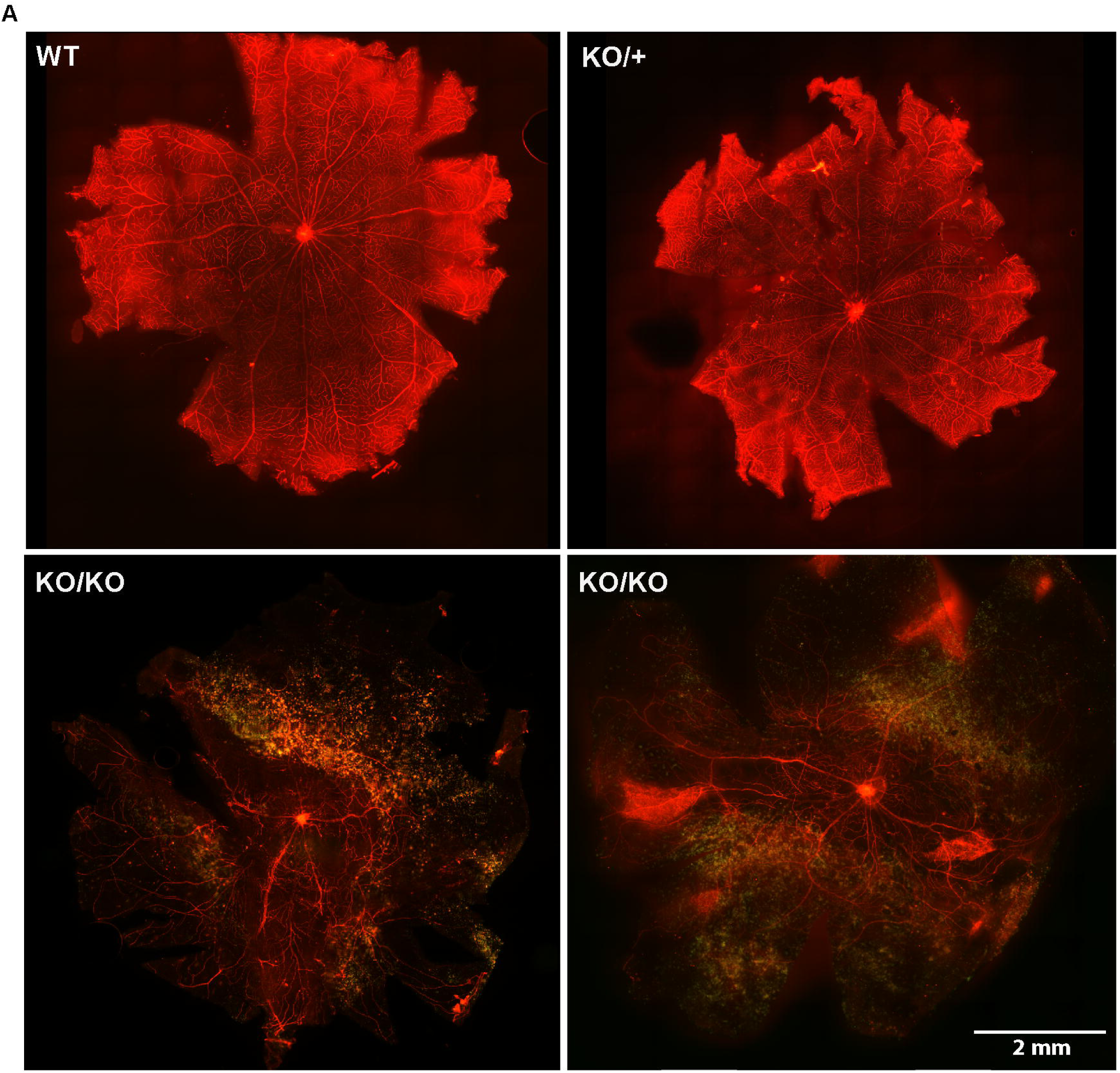
Effect of the 18-bp deletion in 3-month-old *Lrp5*^*KO1*^ rats on vascularization of the retina. Flat-mounted retinas of wild-type (WT) and heterozygous (KO/+) animals stained with Alexa Fluor 594–conjugated isolectin B4 had well-organized blood vessels originating at the optic disc and pervasive branching into the retinal periphery. Vessels in retinas from two *Lrp5*-null knockout rats were sparse and disorganized, with reduced branching, and extensive autofluorescent exudates were present. The images are representative of 8 WT, 5 KO/+, and 6 KO/KO rats.

**Fig. 6.**
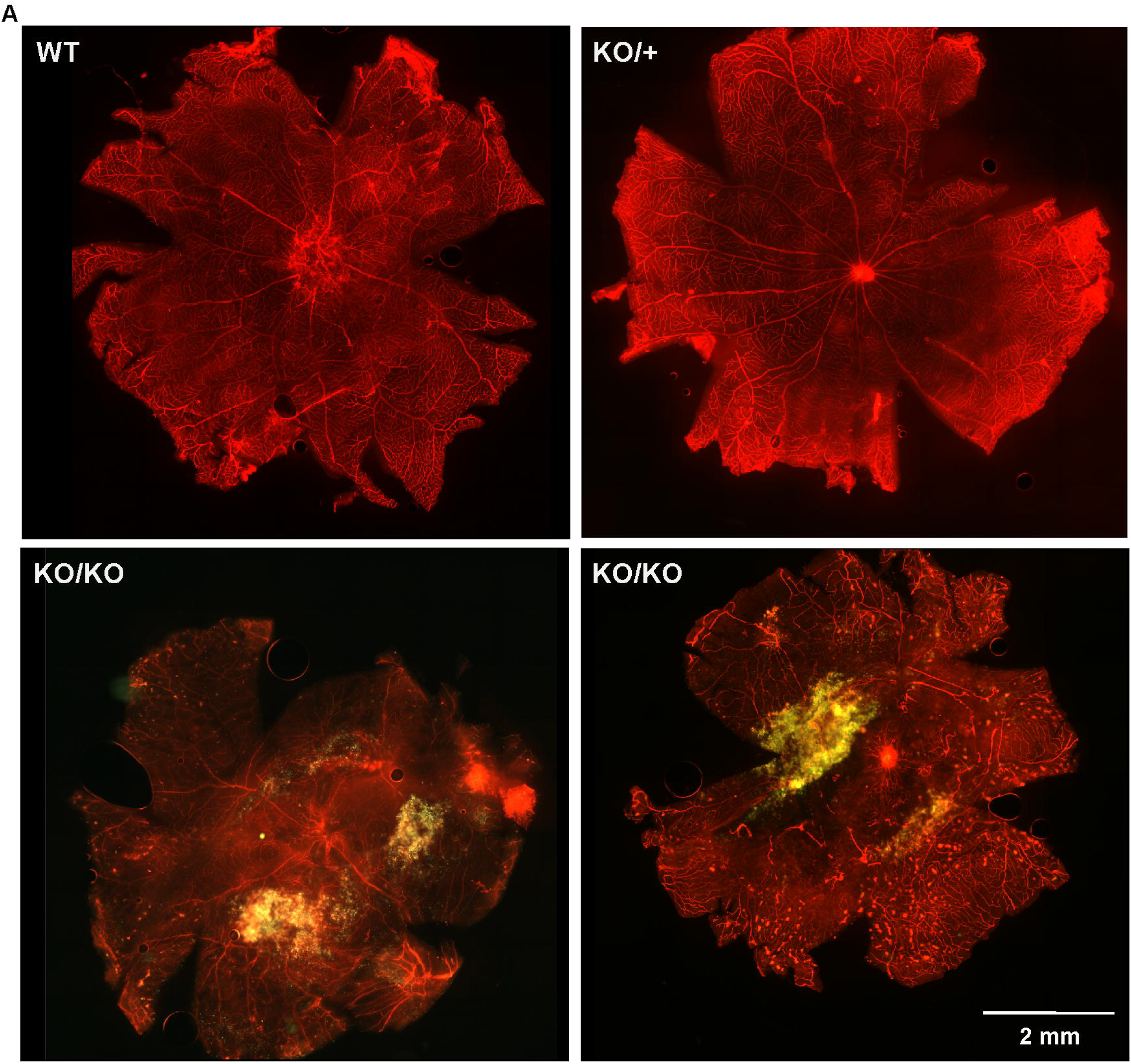
Effect of the inactivation of Lrp5 by allelic inversion on the retinal vascularization of 3-month-old *Lrp5*^*KO3*^ rats. While the vasculature of WT and KO/+ rats was normal, retinal vessels of *Lrp5*-null knockout rats were sparse and disorganized, with reduced branching and vessels ending in vascular tufts (right image), and extensive autofluorescent exudates were present. The images are representative of 6 WT, 7 KO/+, and 8 KO/KO rats.

Three-month-old rats have a completely developed superficial retinal vasculature extending to the periphery, with the extensive, well-organized branching pattern expected in wild-type animals. The retinas of heterozygous rats showed a similar pattern and appeared to be normally developed in animals with either the 18-bp deletion (*Lrp5*^*KO1*^) or the allelic inversion (*Lrp5*^*KO3*^) in only one *Lrp5* alelle (Figs. 5 and 6).

The superficial retinal vasculature of knockout *Lrp5*^*KO1*^, *Lrp5*^*KO*2^, and *Lrp5*^*KO3*^rats had a markedly abnormal phenotype. The number of large vessels radiating from the optic disc was reduced. The vasculature was disorganized, the network of branching, small vessels was nearly absent in large regions of the retina, and in some regions the vessels did not reach the peripheral retina (Figs. 5-7). Some vessels also ended in “glomerular” tufts (Fig. 6, right KO/KO image)

### Quantitative analysis of the retinal vasculature using AngioTool

To verify that loss-of-function mutations in *Lrp5* cause impaired retinal vascularization, the AngioTool image analysis program was used to quantify the vascularized area, vessel length, and branch point density in retinas from *Lrp5*^*KO1*^ and *Lrp5*^*KO3*^ rats. Because of the large regions of exudate. in the retinas of *Lrp5*^*KO2*^ rats (Fig. 7), it was not possible to quantify the vascular pathology in this strain.

**Fig. 7.**
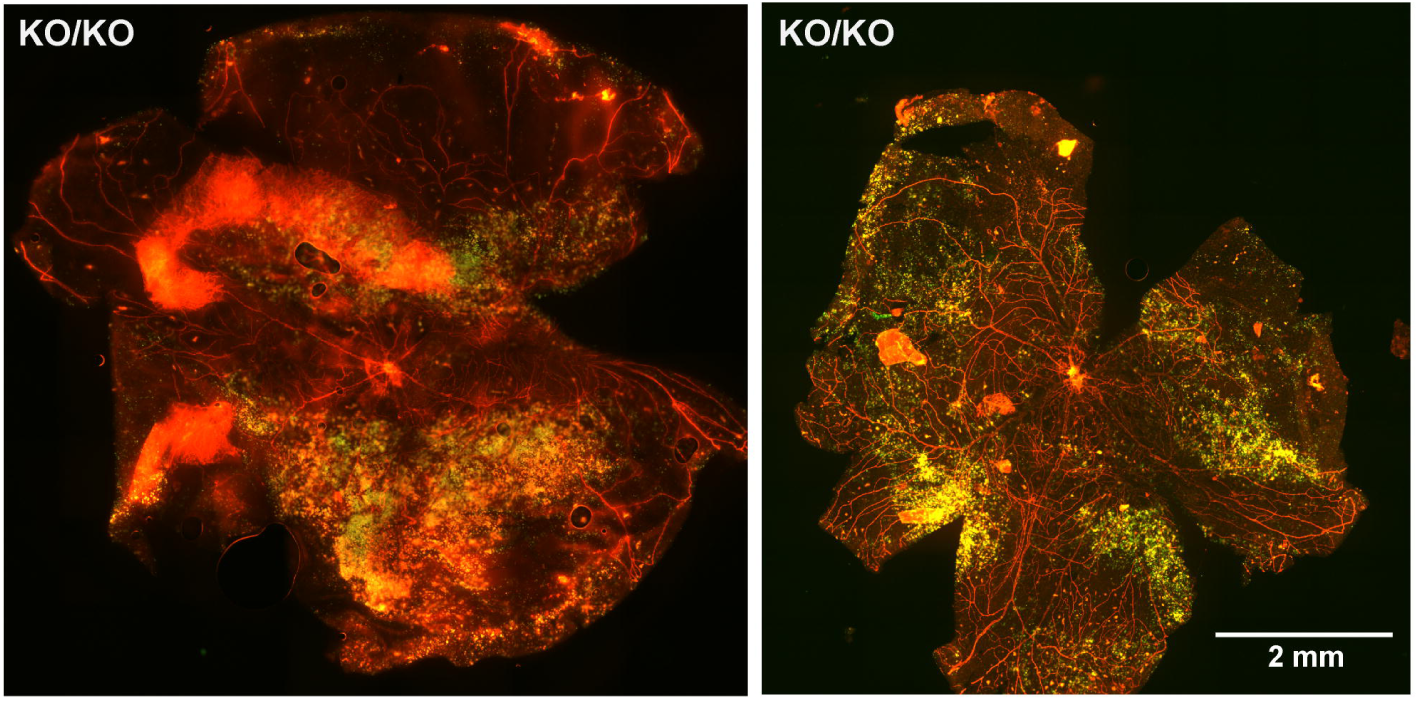
Retinas of 3-month-old, homozygous *Lrp*^*KO2*^ rats that carrried the 22-bp deletion. Exudates and abnormal development of the vasculature are evident. Images are representative of seven rats.

Because the large exudates in the *Lrp5* knockout retinas obscure the vasculature, it was not possible to make measurements of the entire retina. Thus, rather than measuring the vascularized area of the entire retina, as reported in previous studies of young mice with an *Lrp5* mutation [16, 18], in this study, that area is defined as the percentage of retinal area that is vascularized within a 4-mm^2^ digital explant (Fig. 8).. These explants were taken from areas of wild-type, heterozygous, and homozygous animals where the vasculature was fully visible and where there were minimal cuts in the retina that were made in the process of flat mounting. Where possible, two explants were analyzed from the right and left retinas of each animal, although in some retinas it was possible to analyze only one explant.

**Fig. 8.**
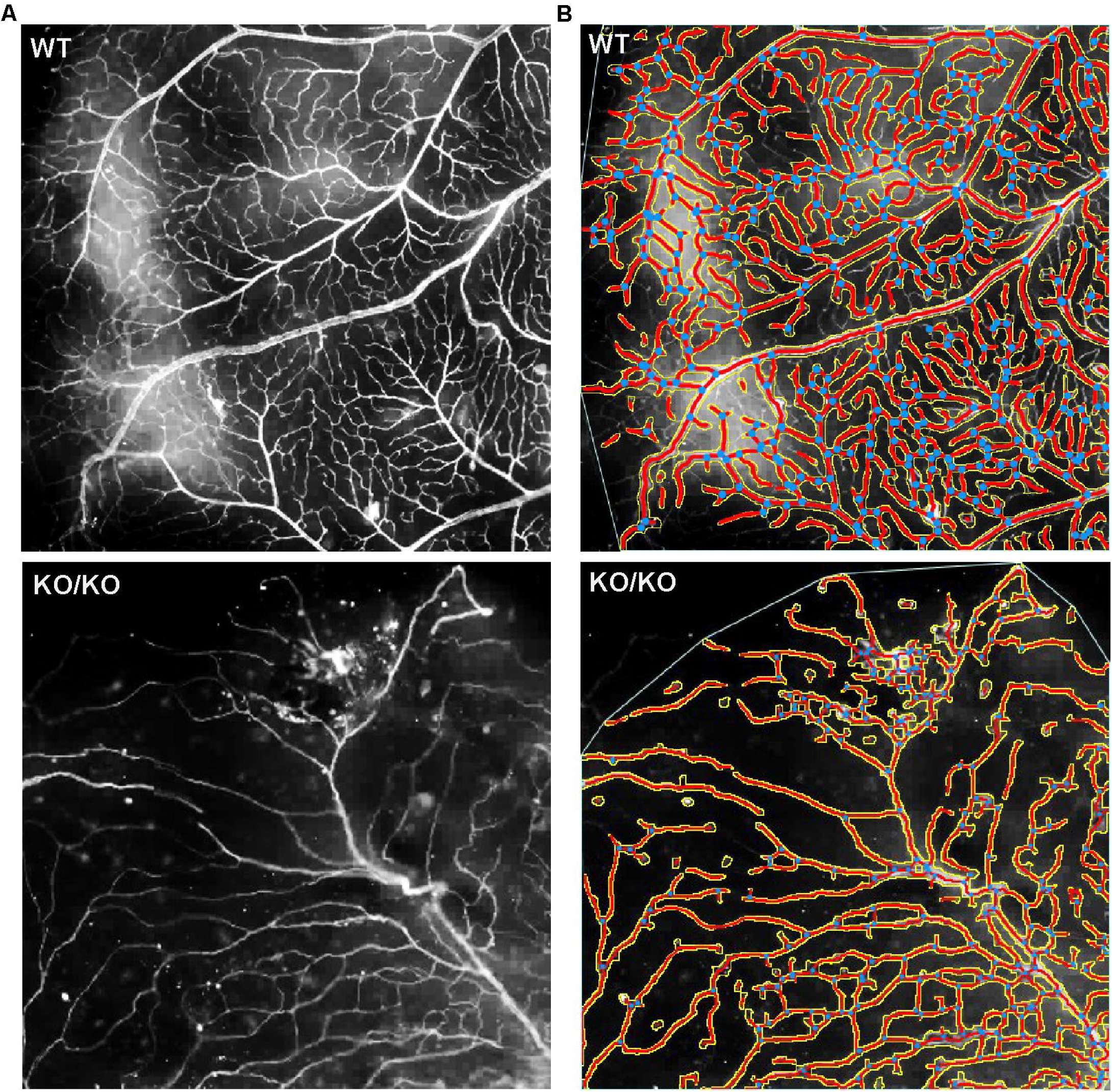
Representative images of 4-mm^2^ digital explants of vascular networks analyzed by AngioTool. Explants of wild-type (WT) and knockout (KO/KO) retinas with the 18-bp deletion (*Lrp5*^*KO*1^) are shown in the left panels. The right panels show Angiotool images with vessels outlined in yellow and highlighted in red, with branch points marked in blue. The program calculates the vascularized area, vessel length, and branch point density.

In *Lrp*^*KO1*^ rats (homozygous 18-bp *Lrp5* deletion), 25.4% of the retina was vascularized, significantly different than the 37.7% vascularization of wild-type retinas (Fig. 9). The median vessel length of wild-type retinas was 1.75 mm, but there was a significant reduction, to 0.63 mm, in *Lrp5*^*KO1*^ retinas (Fig. 9B). The decreased branching of vessels was notable on images of homozygous knockout retinas (see Figs. 5, 6 and 8). Analysis showed that in rats with the 18-bp deletion, there were 86.3 branch points/mm^2^ in wild-type retinas and 42.3/mm^2^ in *Lrp5*^*KO1*^ retinas, a statistically significantt reduction (Fig. 9C). None of the measured parameters in heterozygous *Lrp5*^*KO1*^ retinas were different from those of the wild type.

**Fig. 9.**
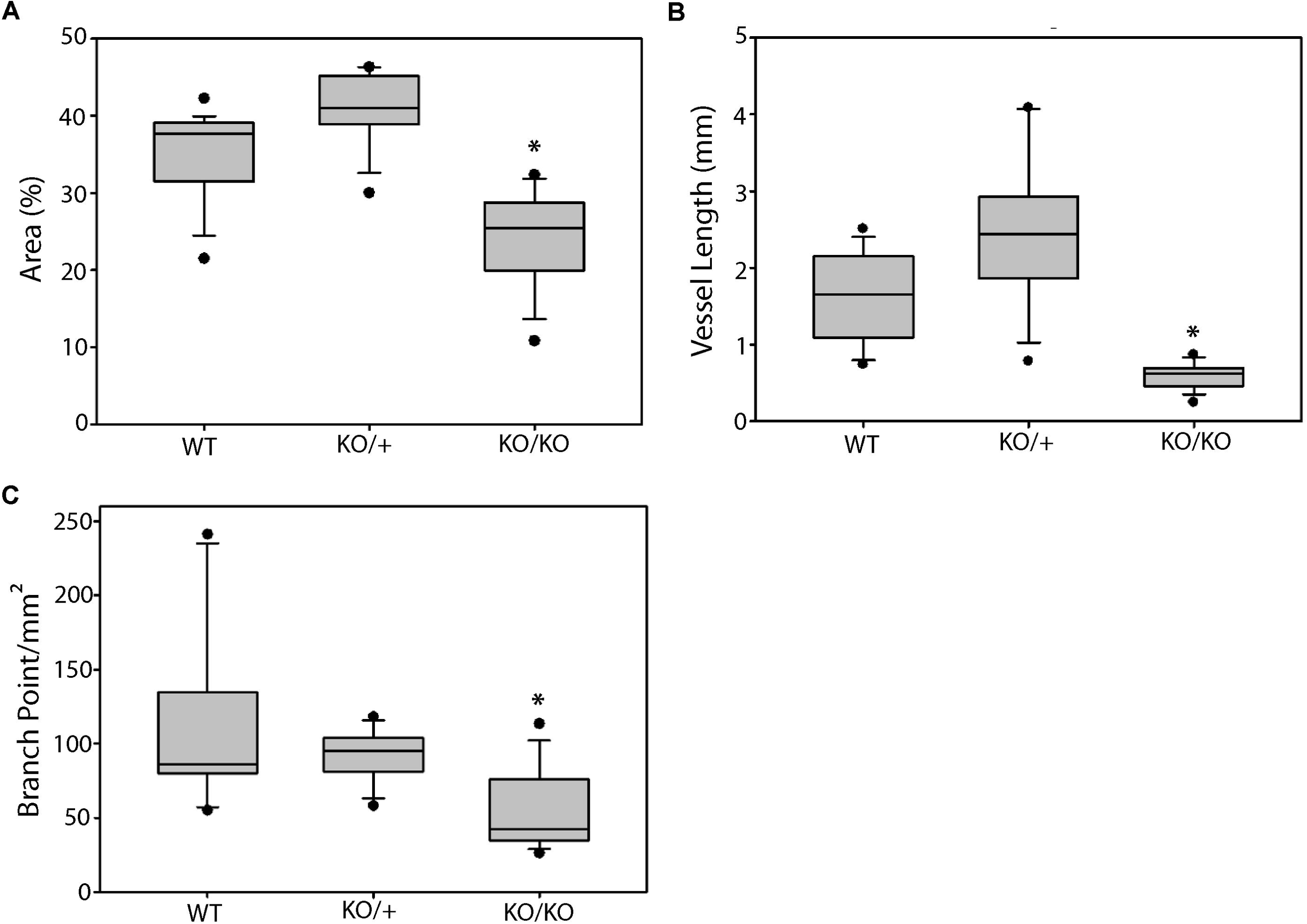
Effect of the 18-bp deletion (*Lrp5*^*KO*1^) on the retinal vasculature. The vascularized area, vessel length, and number of branch points were significantly reduced, while the retinas of heterozygous rats (KO/+) were not significantly different than retinas of wild-type (WT) rats. Kruskal-Wallis one-way analysis of variance on ranks and Dunn’s method. * p < 0.05. Number of explants: WT, *n* = 37; KO/+, *n* = 23; KO/KO, *n* = 31.

The inactivation of *Lrp5* in *Lrp*^*KO*3^ mice (allelic inversion) had a effect on the retinal vasculature similar to that of the 18-bp deletion. The vascularized area was reduced from 40.7% in wild-type rats to 32.7% in *Lrp5*^*KO*3^ rats (Fig. 10A). The median vessel length was reduced from 1.9 mm to 0.9 mm (Fig. 10b), and the median branch point density in the *Lrp5*^*KO3*^ retinas was only 48.8/mm^2^, compared with 97.5/mm^2^ in wild-type retinas (Fig. 10C). All of these differences were statistically significant. As in rats with an 18-bp deletion, the heterozygous *Lrp5*^*KO3*^ genotype had no effect on the retinal vasculature.

**Fig. 10.**
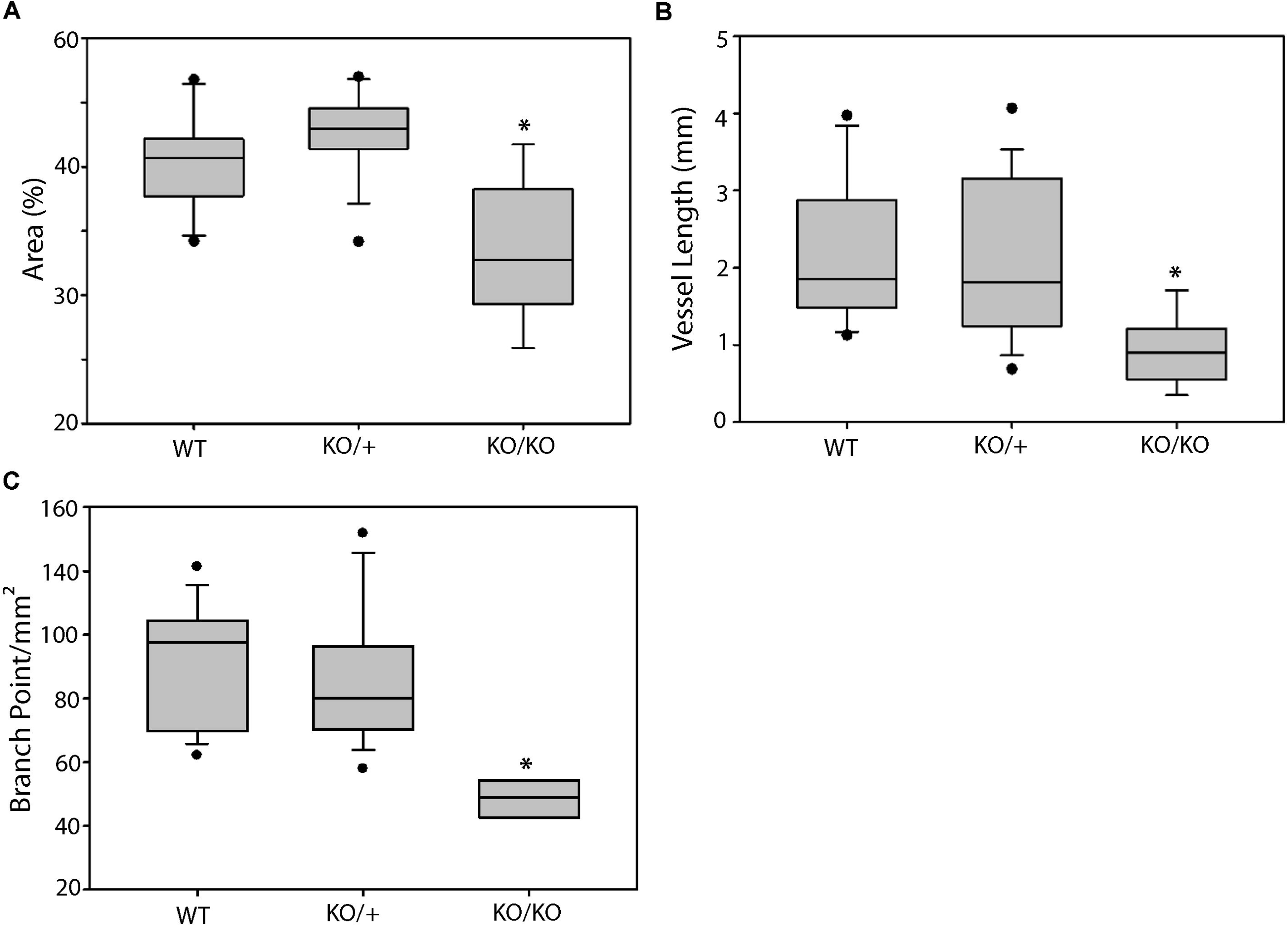
The retinas of *Lrp5*^*KO*3^ rats (allelic inversion) had significantly reduced the vascularized area, vessel length, and number of branch points. Heterozygous (KO/+) retinas were not significantly different than wild-type (WT) retinas. * = p < 0.05. Number of explants: WT, *n* = 29; KO/+, *n* = 37; KO/KO, *n* = 15.

## Discussion

In this study, we produced three strains of rats with loss-of-function mutations in the *Lrp5* gene using CRISPR/Cas9-mediated gene editing. The loss of this co-receptor inhibited Wnt signaling required for normal development of both trabecular and cortical bone. In trabecular bone, this resulted in decreased bone mineral density, bone mass, trabecular thickness, and trabecular separation. Cortical bone density did not change, but bone size was decreased. The *Lrp5* mutations also caused retinal pathology that closely resembled familial exudative vitreoretinopathy. Retinal vascularization was abnormal, with a reduced area of vascularization and reduced branching of vessels. Notably, the animals had extensive retinal exudates.

We have successfully developed the first rat model of osteoporosis by modifying the *Lrp5* gene. The availability of this genetically modified rat increases the options available for evaluating osteoporosis therapies, and the larger size of the rat relative to the mouse may facilitate the assessment of orthopedic procedures in the context of Lrp5 deficiency.

The retinal vascular pathology relative to their control littermates associated with *Lrp5* knockout was similar in all three rat lines and was similar to that of *Lrp5* knockout mice [11, 15], validating this model for studies of the role of Wnt signaling in the control of retinal vascular development. The retinas used in this study were from 3-month-old adult rats and had advanced vascular disease. Most previous studies of *Lrp5* knockout mice concentrated on early postnatal development of the retinal vasculature [11, 15-17]. In two studies that included 1-month-old mice, a reduced vasculature with vessels ending in tufts was also observed [11, 17]. Further, the expression in *Lrp5*-null rats of a sparce, tortuous retinal vasculature with vessels that end in bulbous tufts also suggests that the rat retinal pathology may prove to be a good model of human *Lrp5* mutations that cause FEVR [32, 33].

The *Lrp5* knockout rat retinas had extensive, autofluorescent exudates (see Figs. 5-7), similar to those seen in human FEVR retinas [13, 16, 32,33]. To our knowledge, autofluorescent exudates have not been reported in adult *Lrp5* knockout mouse models, and exudates are not detectable in published images from younger (P7 – P17) *Lrp5* knockout mice [11, 15, 17]. Fluorescein leakage from retinal vessels of *Lrp5* knockout mice has been demonstrated, and retinal hemorrages were occasionally seen on in vivo fundoscopic images of 2.5-month-old *Lrp5*-null mice [18]. In agreement with the exudative pathology seen in humans who have mutations in the Wnt signaling pathway, our data suggest that the knockout of *Lrp5* in rats causes a defective blood–retinal barrier. Diaz-Coranguez et al. have reviewed the literature showing that Wnt signaling and norrin are essential to the development of the blood–retina barrier [34].

## Conclusion

This study demonstrates the feasibility of using CRISPR/Cas9 to produce genetically modified lines of rats. Our model of *Lrp5* mutation in rats will be useful in further studies of Wnt signaling in general. More specifically, the presence of bone and retinal pathology will make these lines useful for research on the treatment of osteoporosis and FEVR. Further studies of the rat retinas, including histology, fundus photography, optical coherence tomography, measurement of vascular permeability and electroretinography, are in progress.

## Acknowledgements

This work was funded by the Van Andel Institute and the Calvin Fund for Eye Research. Key members of the VARI Vivarium and Transgenics Core included Bryn Eagleson, Adam Rapp, Nicholas Getz, Audra Guikema, Tristan Kempston, Malista Powers, and Tina Schumaker. We thank other members of the Williams Laboratory for their advice and assistance, and David Nadziejka for editorial assistance.

## Figure Legends

**Suppl. Fig. 1. *Lrp5*-deficient rats showed a decrease in total femoral BMD.** Aerial BMD of femurs was measured for *Lrp*^*KO1*^, *Lrp5*^*KO2*^, and *Lrp*^*KO3*^ rats using DXA, *n =*4-9. For all graphs, * = *p* < 0.05.

