## Supplementary figures and images for "Low-density Lipoprotein Receptor-related Protein 5 (LRP5)-deficient Rats Have Reduced Bone Mass and Abnormal Development of the Retinal Vasculature"

### Supplemental Figure 1

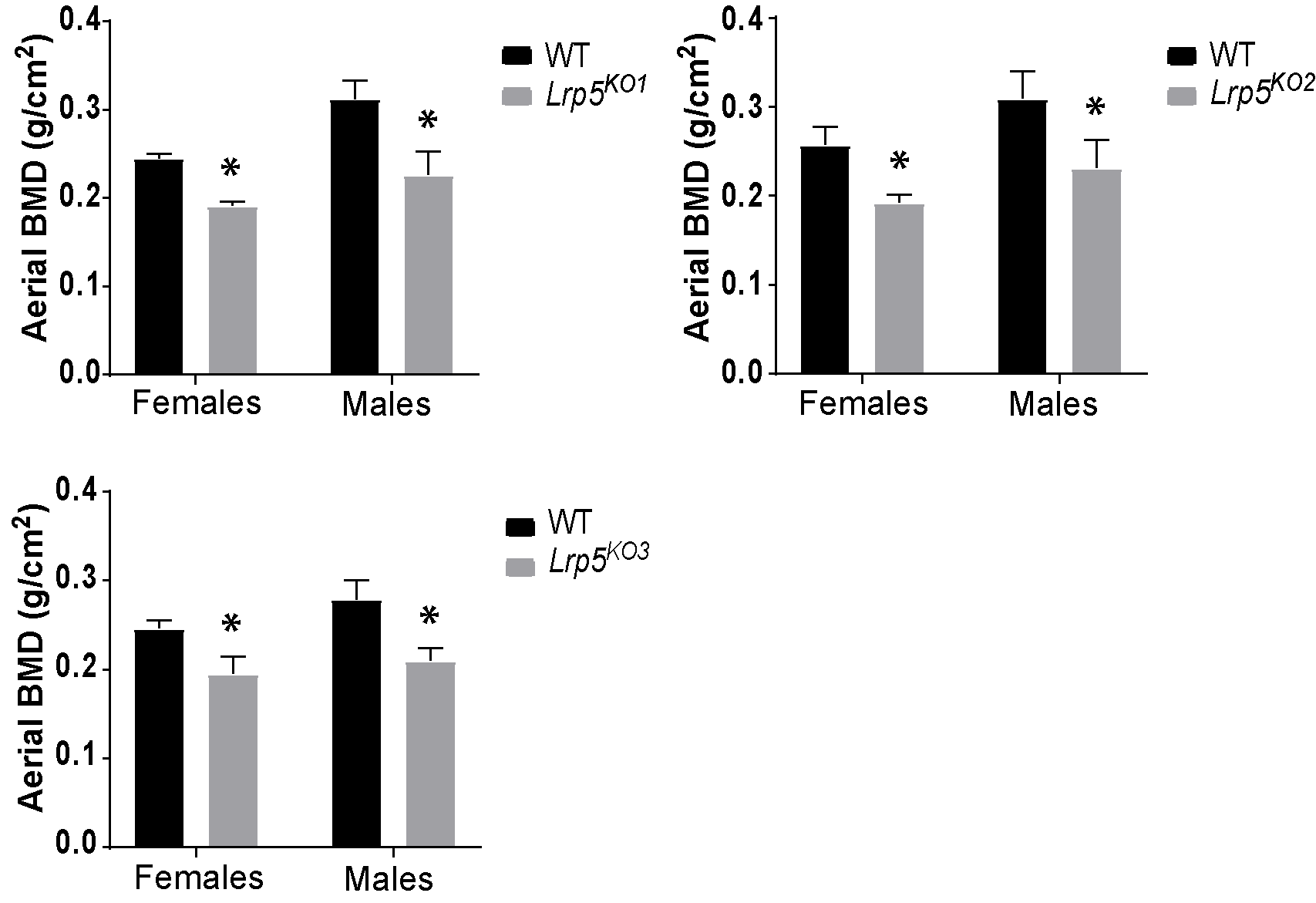
